# Development of the Pre-Gnathal Segments of the Insect Head Indicates They Are Not Serial Homologues of Trunk Segments

**DOI:** 10.1101/2020.09.16.299289

**Authors:** Oren Lev, Ariel D. Chipman

**Affiliations:** The Dept. of Ecology, Evolution & Behavior, The Silberman Institute of Life Sciences, The Hebrew University of Jerusalem

## Abstract

The three anterior-most segments in arthropods contain the ganglia that make up the arthropod brain. These segments, the pre-gnathal segments, are known to exhibit many developmental differences to other segments, believed to reflect their divergent morphology. We have analyzed the expression and function of the genes involved in the segment-polarity network in the pre-gnathal segments compared with the trunk segments in the hemimetabolous insect *Oncopeltus fasciatus*. We show that there are fundamental differences in the way the pre-gnathal segments are generated and patterned, relative to all other segments, and that these differences are general to all arthropods. We argue that given these differences, the pre-gnathal segments should not be considered serially homologous to trunk segments. This realization has important implications for our understanding of the evolution of the arthropod head. We suggest a novel scenario for arthropod head evolution that posits duplication of an ancestral single-segmented head into three descendent segments. This scenario is consistent with what we know of head evolution from the fossil record, and helps reconcile some of the debates about early arthropod evolution.

## Introduction

Arthropods are a hyper-diverse animal phylum characterized by both very high species numbers and exceptional biomass. One of the reasons for their immense success is believed to be their segmented body plan, which provides high evolvability via a modular organization (1).

Segments are repeating body units along the anterior to posterior body axis, which form in a developmental process known as segmentation. Although the specifics of the segmentation process vary among species (2, 3) and can even vary within an individual embryo (4, 5), core aspects of the process are highly conserved, and the segments are generally accepted to be homologous between all arthropods and serially homologous within an individual (6, 7). Segmentation is often described as a hierarchical process, based on what is known from the *Drosophila* segmentation cascade, and the different stages of the process are named after the stages in the *Drosophila* mode. Most relevant to comparative analyses of segmentation are the genes of the two last stages: pair-rule genes and segment-polarity genes (2, 8, 9). The pair-rule genes are expressed in a two-segment periodicity in *Drosophila* and in many other holometabolous insects. Their ancestral role is probably to generate the preliminary repeated pattern that is at the base of segmentation (9). They often interact in a complex gene regulatory network (GRN), the details of which vary among species (10-14). Segment-polarity genes are expressed in segmental stripes in *Drosophila* and in all other arthropods studied to date. Their role seems to be defining segmental (or parasegmental) boundaries, and they function as a highly conserved GRN with very similar interactions among and within all studied arthropods (2, 15, 16). The segment-polarity GRN is based upon the interactions of two signaling pathways, Hedgehog and Wnt, and the paralogous transcription factors Engrailed and Invected. Later identity of the segments is conferred by Hox genes, probably together with other genes (17).

During development, segments are grouped into functional body units called tagmata. The organization of segments into tagmata varies among different arthropod clades, e.g. insects have three tagmata: head, thorax, and abdomen, while chelicerates have two: prosoma and opisthosoma. There is evidence to suggest that the differences between tagmata may have an early developmental basis (1).

The embryonic insect head can be divided into two areas: the anterior procephalon and the posterior gnathocephalon, which can be clearly distinguished based on their shape during the germband stage. The developing segments of the gnathocephalon are very similar in size and shape to the thoracic segments that lie posterior to them. They develop to give rise to the three segments that carry the adult mouthparts: the mandibles, maxillae and labium (18). The embryonic procephalon makes up the head-lobes, an enlarged region without clearly distinguishable segments, which develops to give rise to three adult segments: the ocular, antennal and intercalary segments, each of which has a dorsal ganglion. The three ganglia together make up the tri-partite brain. Because these three segments lie anterior to the limb-bearing gnathal segments in insects, we refer to them from here on collectively as the pre-gnathal segments (PGS). For simplicity, we will refer to all segments posterior to the PGS as trunk segments. There is mounting evidence that the three anterior segments in all arthropods are homologous and share a deep evolutionary history (19). When discussing them in a general arthropod comparative framework, they are usually referred to by the names of the homologous ganglia they carry, namely (from anterior to posterior) the protocerebral, deutocerebral and tritocerebral segments.

It has long been known that the PGS are developmentally distinct from other segments (20). One significant difference between the PGS and the rest of the body segments is that pair-rule genes are not expressed in them during early development (18, 20). The later acting Hox genes, which are involved in conferring morphological identity to specific segments, are only expressed in the tritocerebral segment and not in the anterior two PGS.

The segment polarity genes are expressed in the PGS, but with some variations relative to what is found in other segments. Most notably, the timing of segment polarity gene expression is different: e.g., the segment polarity gene *engrailed* (*en*) is first expressed in the mandibular segment even though the PGS are already patterned at that stage (5, 21, 22). Furthermore, in several cases, *hedgehog* (*hh*) is expressed as a single domain that splits once or twice to give rise to three stripes, each marking a region of a different one of the anterior segments. In the spider *Parasteatoda tepidariorum* and the centipede *Strigamia maritima, hh* expression splits twice, from the anterior stripe or the posterior stripe respectively (23-25). In the millipede *Glomeris marginata* (26), the beetle *Tribolium castaneum* (27) and the cricket *Gryllus bimaculatus* (28), *hh* expression splits only once, with a third expression domain appearing later, marking the tritocerebral segment. Work in the model species *Drosophila melanogaster* showed unexpected interactions and expression domains of the different segment polarity genes in the anterior head segments (29). The authors of this work focused on relatively late expression and interpreted the differences relative to other segments as being related to the different morphology of the structure formed from these segments.

While these previous studies have pointed out several unusual characteristics of the patterning of the pre-gnathal segments, there is no single species for which there exist both descriptive and functional studies in both early and late stages of the formation of the PGS. Furthermore, no previous study has attempted to reconstruct the sequence of gene interactions involved in generating these segments and to compare this sequence to that found in other segments. We aim to fill this gap, using the milkweed bug *Oncopeltus fasciatus* as our chosen study organism for this analysis. Unlike *Drosophila*, components of the larval head are generated in full during early development of *Oncopeltus*, making it possible to study the development of all head structures in the embryo (30). The earliest stages of PGS patterning occur during blastoderm segmentation, which has been studied extensively in *Oncopeltus*, is easy to access experimentally and is fairly well-understood (5).

In this paper, we provide a detailed descriptive and functional analysis of the early stages of the formation of the PGS in *Oncopeltus*. We go beyond this basic description, including published data from other arthropod species and linking the developmental results with what is known about the early fossil record of the arthropod head, to build a comprehensive evolutionary-developmental scenario for the evolution of the arthropod head. We argue that the differences in the early determination of the PGS are due to their having a separate origin and a separate evolutionary history from all other segments.

## Results

### Early wildtype expression of components of the segment-polarity gene network

We started by looking at the expression of three segment polarity genes from the earliest stages of formation of the PGS and up to the early segmented germband stage (Fig. 1). The transcription factor encoding gene *invected* is not expressed in the PGS during the blastoderm stage (5). Its earliest expression is after the formation of the head lobes in the germband. Expression of *hh* begins in the mid blastoderm, approximately 30-32 hours after egg laying (hAEL at 25°C). It is first expressed as a ring at the anterior third of the embryo (Fig 1A). Just before blastoderm invagination begins (~36 hAEL) there are two distinct dynamics: a stripe splits from the anterior ring and six more stripes appear more or less simultaneously between the first ring and the invagination site (Fig. 1B. Supp. Fig 1).

**Figure 1.**
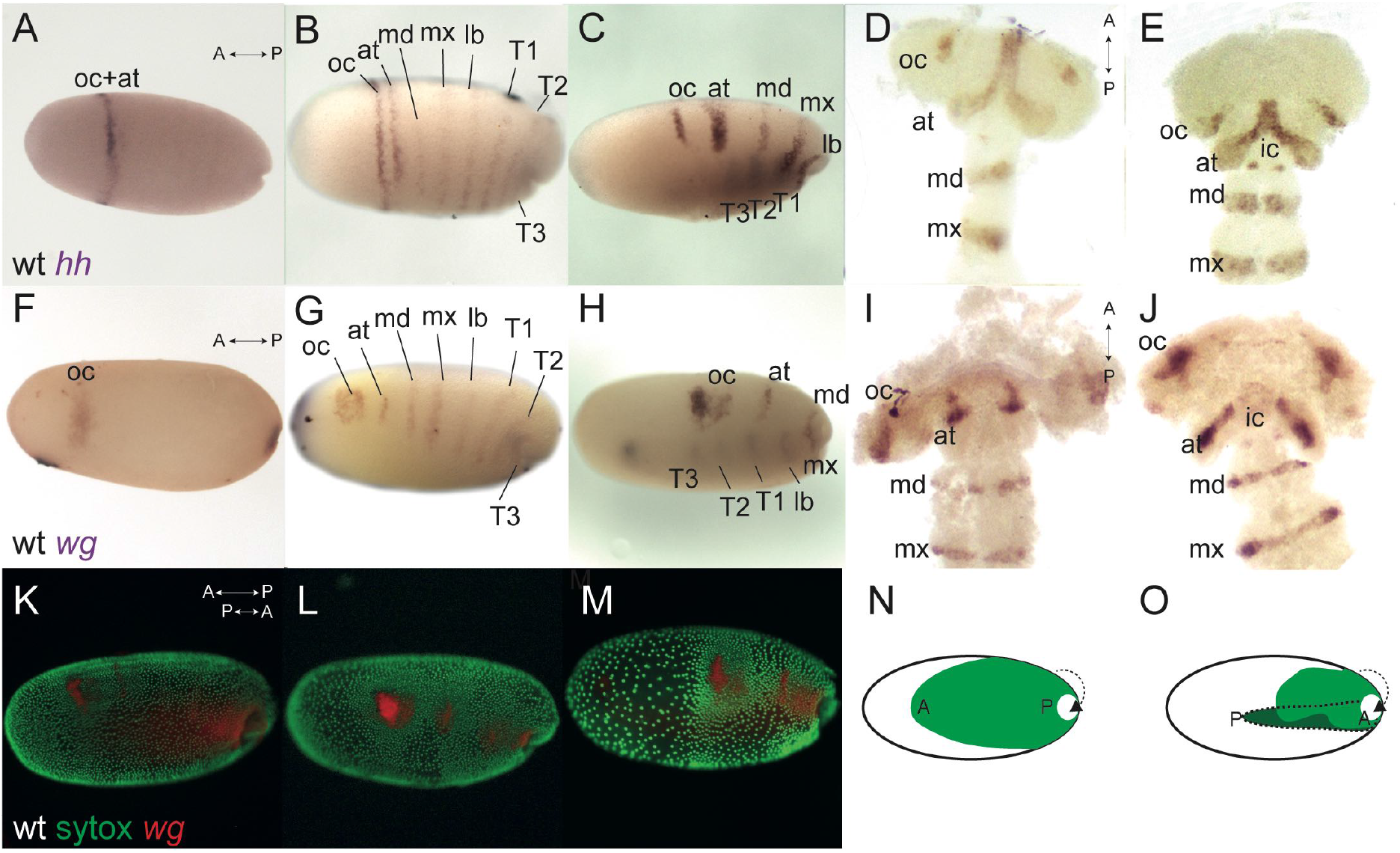
Wildtype expression of *hedgehog* (*hh*) and *wingless* (*wg*) in *Oncopeltus*, before and after blastoderm invagination. A) Expression of *hh* at 32 hours after egg laying (hAEL), late blastoderm stage, is as a ring in the anterior third of the embryo. B) at around 36 hAEL, the *hh* anterior expression ring “splits” into two stripes corresponding to the presumed ocular and antennal segments. six additional stripes of expression appear, corresponding to the presumed gnathal and thoracic segments. C) embryo at mid-invagination, ~42 hAEL. The two anterior stripes are fully separated. Thoracic segments have already invaginated and expression in them can be seen through the yolk. D-E) Germband embryos, ~48-52 hAEL. In the later germband embryo, ~52 hAEL (E), two dots of *hh* expression appear in the presumed intercalary segment, which are not seen in the earlier embryo (D). F) Expression of *wg* at ~32 hAEL is located to an anterior stain and a posterior stain marking the future invagination site. G) at ~36 hAEL 7 stripes of expression appear simultaneously, corresponding to the presumed antennal, gnathal, and thoracic segments. H) embryo at mid-invagination, ~42 hAEL. I-J) Germband embryos, ~48-52 hAEL. In the later germband embryo, ~52 hAEL (J), two dots of *wg* expression appear in the presumed intercalary segment, which cannot be seen in the earlier embryo (I). K-M) The process of invagination from ~36-42 hAEL. Nuclei stained by SYTOX in green and wildtype *wg* expression in red. At this stage, the anterior-posterior axis of the egg is opposite to that of the invaginating embryo. N-O) Illustration of the invagination process. ‘A’ marks the embryonic anterior and ‘P’ the posterior. The invaginated germband is shown in darker green. Abbreviations: at – antennal segment. ic – intercalary segment. lb – labial segment. oc – ocular segment. md – mandibular segment. mx – maxillary segment. T1-T3 – First to third thoracic segments.

Throughout invagination (~36-48 hAEL) the expressions stripes of *Of-hh* become narrower and dorsalized as the two anterior-most stripes fully separate (Fig. 1C-D). Around 50 hAEL there is a new expression of *Of-hh* in two dots (Fig. 1E). Based on their relative position and shape, and based on previous work on blastoderm segmentation (5) we interpret the two stripes that emerge from the splitting of the initial ring as representing the ocular and antennal segments, the six simultaneous stripes as representing the three gnathal and three thoracic segments, and the two late dots as the intercalary segment.

Blastodermal expression of *wg* has been previously described in *Oncopeltus* (5). However in previous work we did not follow expression into the germband stage, and we believe we therefore did not identify all of the expression domains correctly. We thus describe it again here. The earliest expression of *wg* appears in two domains: a posterior spot at the future invagination site, and an anterior irregular domain (Fig. 1F). The anterior domain is slightly anterior and ventral to the *hh* ring that appears at the same time (Fig. 1A). This domain (referred to by some authors as the “head blob” (31)) expands and develops two lobes to give a heart shape (Fig. 1G-H). Contrary to what we claimed in (5), the two lobes never separate completely, and the heart shape continues until the germband stage, where it is found in the area of the head lobes corresponding to the ocular segment (Fig. 1I-J). At ~32-36 hAEL, seven blastodermal segmental stripes appear simultaneously (Fig. 1G). Six of these extend from the dorsal side of the embryo almost to the ventral side, and correspond to the gnathal and thoracic segments. The seventh anteriormost stripe is shorter and corresponds to the antennal segment. This stripe was previously incorrectly identified as the intercalary stripe. In fact, expression of *wg* in the intercalary segment appears only in the germband, as two dots (Fig. 1J), similar to the intercalary expression of *hh* (Fig. 1E).

The shortening and thickening of the segmental stripes of *hh* and *wg* are due to the condensation of the head lobes that occurs during blastoderm invagination, as evidenced by staining for nuclei together with gene expression (Fig. 1K-M). This condensation is presented schematically in Fig. 1N-O.

### Functions of the segment polarity components in the pre-gnathal segments

To test the function of the Hedgehog and Wnt pathways in the anterior of the embryos, we used parental RNA interference (pRNAi) against positive and negative regulators of each pathway. A positive regulator will decrease the pathway’s activity when knocked-down, while a negative regulator will increase it (Table 1). For the Hedgehog pathway, we chose *hedgehog* (*hh*) – encoding the single arthropod ligand that activates the pathway – and *patched* (*ptc*) – encoding the receptor, which downregulates Hedgehog-signaling when active. For the Wnt pathway, we chose *disheveled* (*dsh*) – which encodes an activating intracellular component of the pathway – and *shaggy* (*sgg*) – which encodes an enzyme that phosphorylates the second messenger β-catenin, thus leading to its degradation. We chose to not directly target either the ligand or the receptor in Wnt-signaling, since both have several copies and are at least partially redundant (32, 33).

**Table 1.** Summary of the RNAi experiments

| Signaling network | Hedgehog |  | Wnt |  |
| --- | --- | --- | --- | --- |
|  | Negative (pRNAi-*ptc*) | Positive (pRNAi-*hh*) | Negative (pRNAi-*sgg*) | Positive (pRNAi-*dsh*) |
| PGS | Enlarged | Reduced | Reduced | Disturbed |
| Trunk | Irregular segment boundaries | Irregular segment boundaries | Dorsal-Ventral axis disruptions | Reduced thorax and abdomen |

Reducing Hedgehog activity via *hh*^RNAi^ results in a subtle but consistent phenotype (Fig. 2) - the reduction of PGS-related structures: The eyes are smaller, and the labrum and antennae are reduced or not developed at all (Fig. 2C-D. Compare with Fig. 2A-B). Without the labrum, the hatchling mandibles and maxillae are exposed. The reduction is also visible when comparing WT *wg* expression in the head lobes (Fig. 2E) to the *wg* expression of *hh*^RNAi^ germbands (Fig. 2F). The segmental *wg* stripe representing the antennal segment is weak or nonexistent in the head lobes during the germband stage, and the head lobes are misshapen and missing the very anterior region. The absence of the antennal *wg* stripe is clear from the first appearance of segmental stripes in the mid-blastoderm (Fig. 2H. Compare with Fig. 2G). There is no obvious effect in other regions of the body.

**Figure 2.**
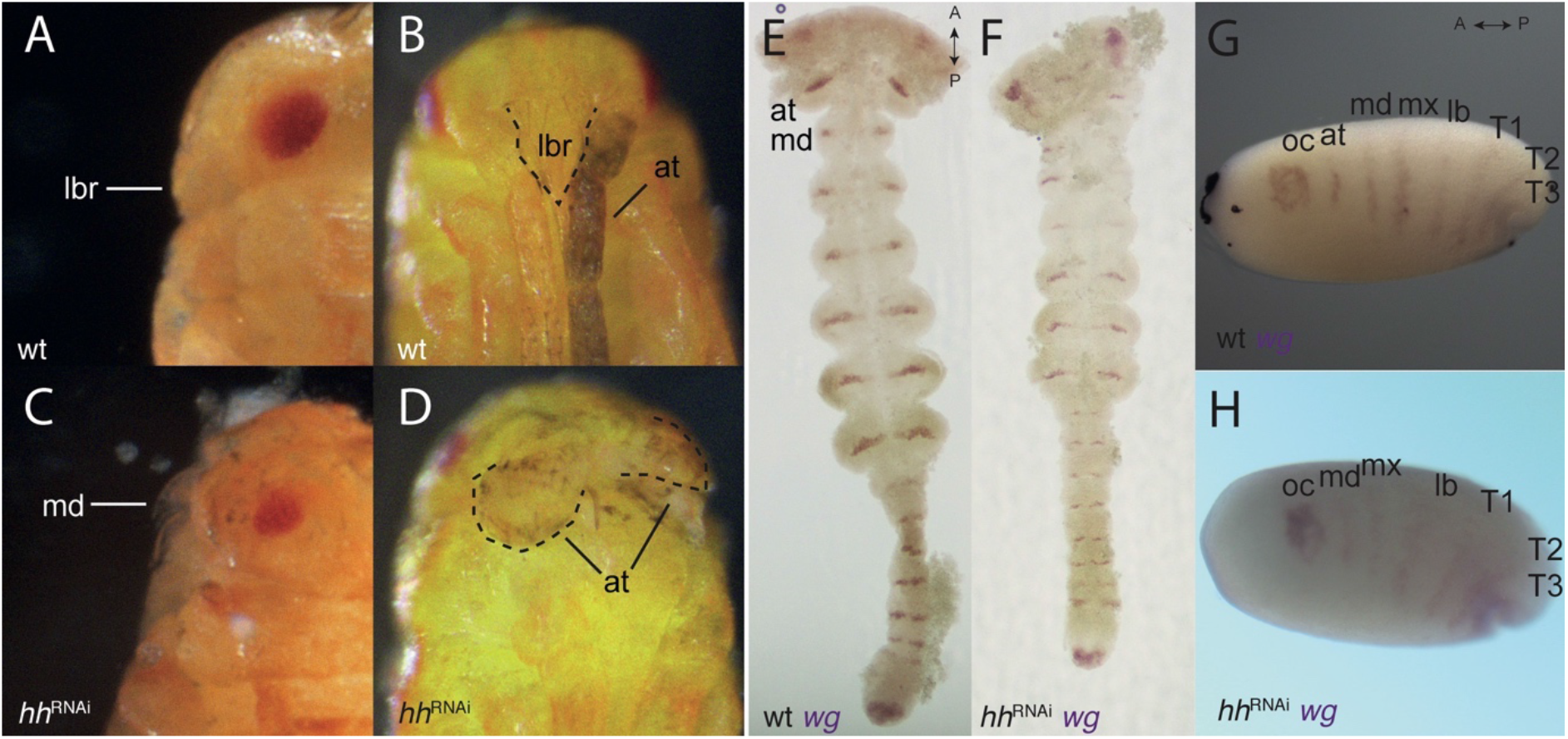
*hh*-RNAi phenotypes. A-B) side and front view of a wildtype hatchling, the labrum covers the mandibles and the antennae lie to its sides. C) a side view of a hatchling with a reduced labrum, exposing the mandibles following a *hh*-RNAi. D) front view of the same hatchling as (C) showing malformed stubby antennae. E) wildtype germband expression of *wg* in the mid germband stage. F) *hh*-RNAi germband expression of *wg*, the antennal expression missing. G) wildtype expression of *wg* in the blastoderm (the same embryo from Fig. 1G). H) *hh*-RNAi expression of *wg* in the blastoderm, the antennal segmental stripe is missing. Abbreviations as in Fig. 1.

Over activating Hedgehog signaling through *ptc* knock-down has a very different effect to *hh*^RNAi^, causing deformation of the trunk segments, but without any clear effect on the head (Fig. 3). This is evident both in knock-down hatchlings (Fig. 3A. Compare with Fig. 3B) and in mounted germbands stained for *wg* (Fig. 3D. Compare with Fig. 3C). Prior to mounting and flattening the germband to a slide, the embryos were far more twisted than are WT germbands (not shown). The blastodermal expression of *wg* is not affected either by over-activating or by reducing Hedgehog signaling (not shown).

**Figure 3.**
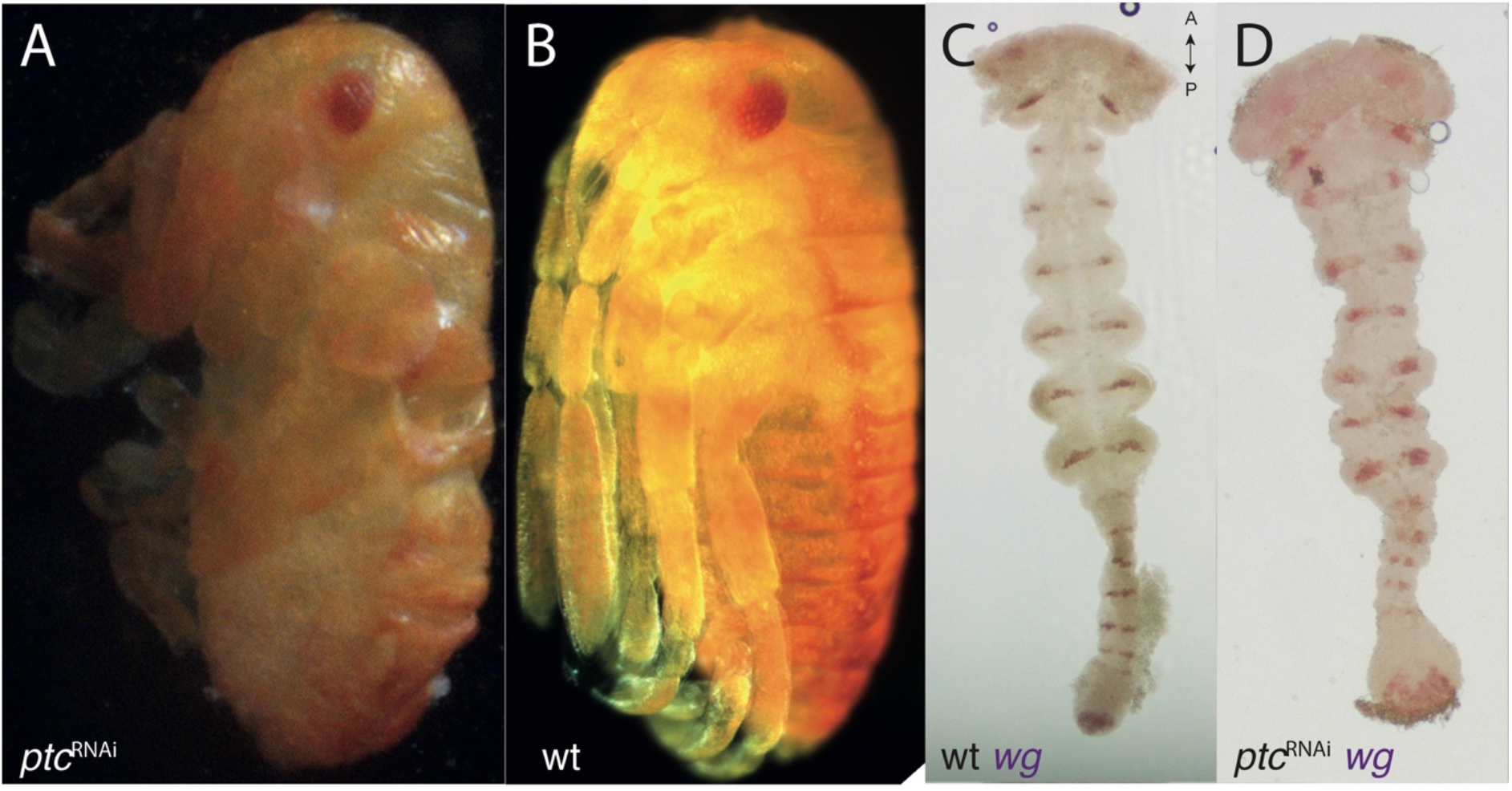
*ptc*-RNAi phenotypes. A) *ptc*-RNAi hatchling with segmental boundaries and limb deformities compared to B) wildtype hatchling of the same age. C) wildtype expression of *wg* in the germband (same embryo from Fig. 2E). D) *ptc*-RNAi expression of *wg* in the germband showing a normal head and mis-formed trunk segments.

When reducing the activity of the Wnt pathway via *sgg*^RNAi^ (fig. 4) and increasing it via *dsh*^RNAi^ (Fig. 5), we see opposite and complementary effects. We find that *sgg*^RNAi^ hatchlings have a severely reduced head (Fig. 4D). Because the head region is missing in these embryos, and the head is normally the last structure to invaginate (Fig. 1L-M) they do not complete invagination properly. We were thus unable to dissect germband stage *sgg*^RNAi^ embryos. Whole mount images of these embryos, show no evidence of anterior expression of *wg* (Fig. 4B). The early expression of *hh* in *sgg*^RNAi^ embryos is diffuse and broader than normal in some embryos (Fig. 4C), whereas in others it is completely disrupted (Fig. 4C’).

**Figure 4.**
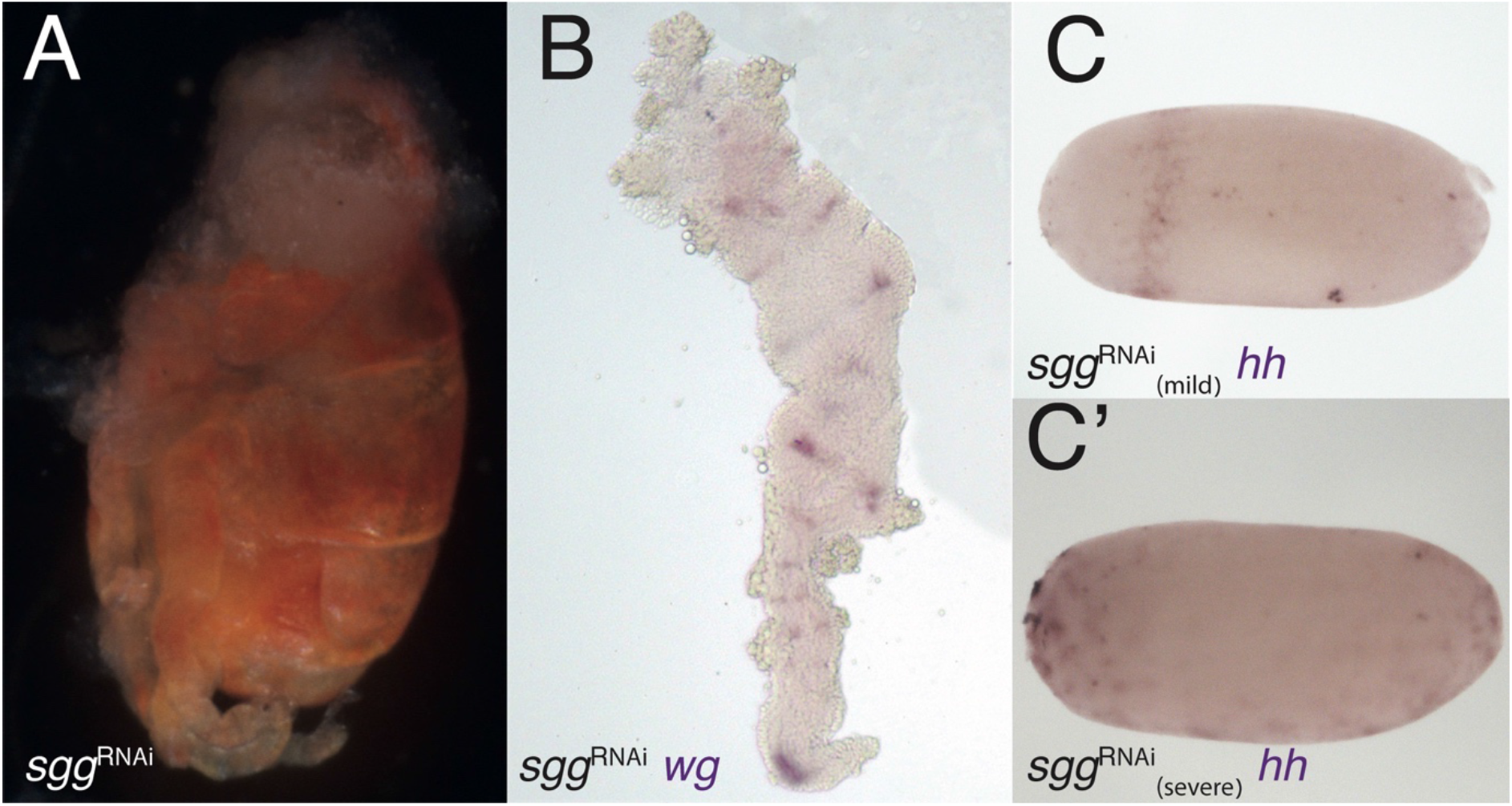
*sgg*-RNAi phenotypes. A) *sgg*-RNAi hatchling with no head and a reduced abdomen (Compare with Fig. 3B). B) Expression of *wg* in *sgg*-RNAi germband embryo. The germband is severely deformed, with no head lobes and no pre-gnathal segments. The exact identity of the remaining segments is difficult to determine. C) Expression of *hh* in *sgg*-RNAi blastoderm embryo with mild knock-down effect. The expression ring is jagged and not uniform. C’) Expression of *hh* in *sgg*-RNAi blastoderm embryo with severe knock-down effect showing patchy expression suggesting that the embryo did not develop at all.

**Figure 5.**
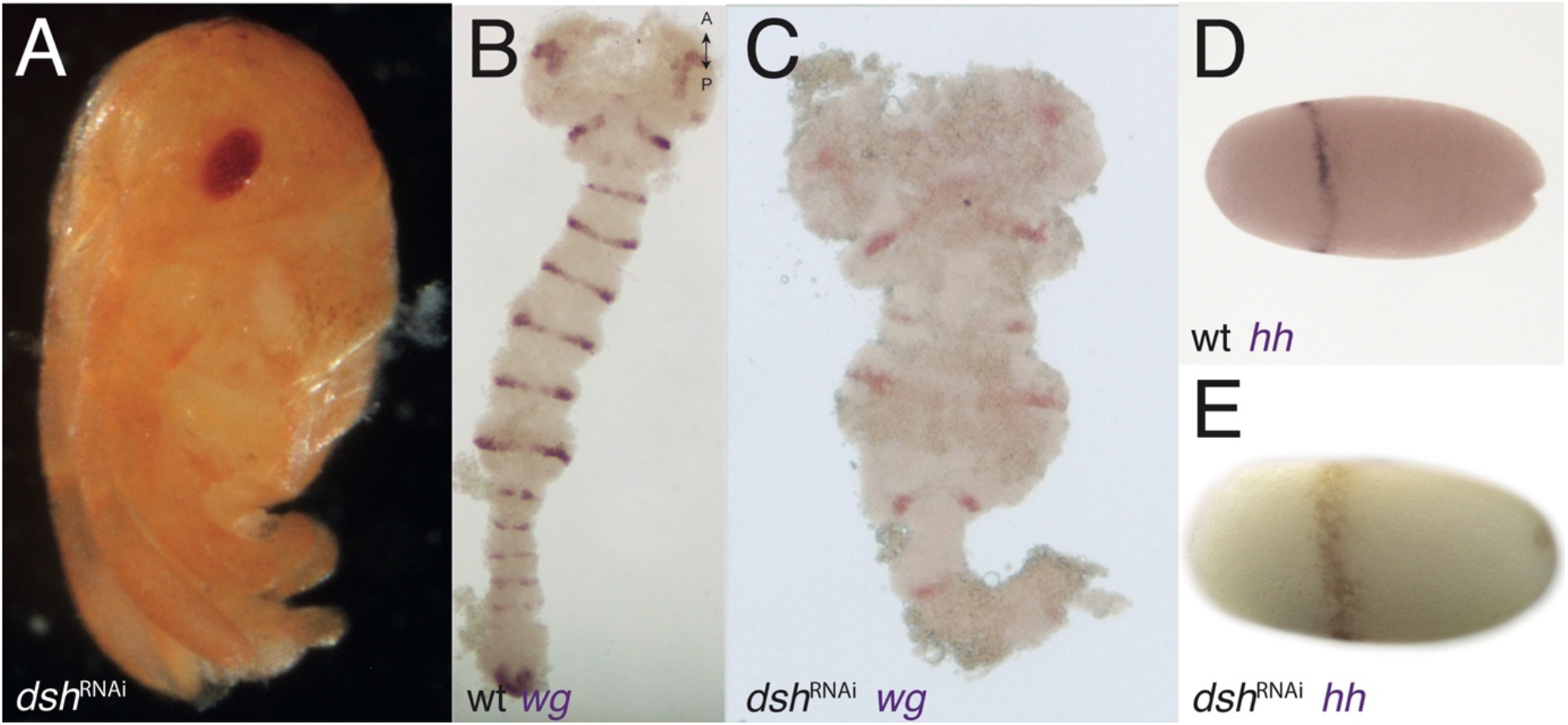
*dsh*-RNAi phenotypes. A) *dsh*-RNAi hatchling with no abdomen (Compare with Fig. 3B). B) Expression of *wg* in wildtype germband stage embryo (slightly older than the one in Fig. 2E). C) Expression of *wg* in *dsh*-RNAi germband with normal PGS but deformed gnathal segments and missing thoracic and abdominal segments. D) Expression of *hh* in wildtype blastoderm stage embryo (Same embryos from Fig. 1A). (E) expression of *hh* in *dsh*-RNAi blastoderm embryo, the anterior ring is jagged and not uniform.

In contrast, *dsh*^RNAi^ hatchlings have a reduced abdomen or, in more severe phenotypes, a reduced abdomen and thorax (Fig. 5A). and severely disrupted and truncated germbands, with almost normal head lobes in *dsh*^RNAi^ embryos (Fig. 5C. Compare with 5B). The early expression of *hh* in *dsh*^RNAi^ embryos is broader than normal and is shifted posteriorly compared with WT embryos (Fig. 5E. Compare with Fig. 5D). Early expression of *wg* in embryos with the Wnt pathway disrupted also have complementary expression patterns. In *sgg*^RNAi^ embryos the anterior expression patch (the “head blob”) is missing, whereas in *dsh*^RNAi^ embryos, the posterior patch is missing (Supp. Fig. 2).

## Discussion

### The expression of segment polarity genes in the PGS of *Oncopeltus* is different from all other segments

Previous studies on segment polarity gene expression in *Oncopeltus* (4, 5) identified two distinct dynamic modes for these genes: segmental stripes of expression of segment-polarity genes appear simultaneously in the gnathal and thoracic segments, and sequentially from a segment addition zone in the abdominal segments. In both cases, *hh* and *inv* are expressed in segmental stripes at the posterior of each segment and *wg* is expressed anterior to them. This expression is the same as that seen in all arthropods studied to date, and presumably reflects the ancient and highly conserved segment-polarity GRN (9, 16, 34).

Conversely, as we show here, the expression of the segment polarity genes is different in the PGS. The expression domains of *hh* and *wg* in the ocular segment are very different in shape and extent, and clearly do not overlap. The extent and pattern of expression of both these genes in the ocular segment are also different from those of all other segments. Expression of these two genes in the antennal segment is similar, although we were unable to determine whether they overlap. Both genes are expressed in a clear stripe, as in other segments.

Expression in the intercalary segment is not in segmental stripes but in two small dots, and these appear very late in development, after all head and thoracic segments are determined and have begun to differentiate. The dynamics of *hh* expression in the PGS are unique, relative to other segment polarity genes in both the PGS and in the rest of the embryo, and involve a stage of a single broad expression stripe splitting to give two stripes. The expression of the best studied segment polarity gene *inv* (an *en* paralog) in the PGS is significantly later than its expression in trunk segments, first appearing only in the germband stage.

### The structure of the segment-polarity network in the PGS in *Oncopeltus* is different to all other segments

Focusing on the functions of the three aforementioned segment polarity genes and on the interactions among them also identifies key differences between what happens in the PGS and what happens in trunk segments. The expression and function of *inv* in *Oncopeltus* has been previously described (although it was originally misidentified as *en*). Knocking down *inv* leads to trunk segments that are moderately or severely disrupted, while the PGS are unaffected (35) indicating a difference in the role of this gene between the two groups of segments.

In *hh*^RNAi^ embryos we find reduced eyes, antennae, and labrum, all of which are structures related to the PGS. However, the “classic” segment polarity knock-down phenotype – disruptions to segment boundaries – is not evident. This is most likely because *hh* is induced by *en* in the trunk segments and is part of a regulatory feedback loop. Therefore, a single knockdown may not result in a noticeable effect, as En/Inv can rescue Hedgehog function through compensation. Because we do see a phenotype in the PGS, we suggest that the segment polarity GRN that maintains *hh* expression in the trunk segments is not active in the same way in the PGS.

This idea is strengthened by *ptc*^RNAi^ embryos. The knock down of *ptc* causes over-activation of the pathway and disrupts normal segmental boundaries in the trunk segments only. This is evident in *wg* stained germbands and in the knock-down hatchlings.

In both *dsh*^RNAi^ and *sgg*^RNAi^ embryos we see irregularities in the expression of *Of-hh* during PGS formation. However, only *sgg*^RNAi^ shows a reduction in the head. This is due to the Wnt pathway’s early function in anterior-posterior axis determination of the embryo (5, 36). Activation of Wnt in the posterior early in bilaterian embryogenesis is crucial for proper pole definition. This process is much earlier than PGS formation, therefore the reduction of the head in *sgg*^RNAi^ is probably a result of a posteriorization of the entire embryo following Wnt over-activation. Conversely, *dsh*^RNAi^ embryos show reduction in the posterior pole of the embryo. Mild cases lack the abdomen, while more severe cases lack all segments up until the gnathal segments. In these embryos the head is not affected by the disruption of the axis. However, the gnathal segments show segmental abnormalities and disrupted *wg* expression, while the PGS are normal. This again suggests that the classical segment polarity GRN does not function in the same manner in the PGS.

### The unique characteristics of the PGS are probably general to all arthropods

Detailed functional studies as we present here have not been done in many other species. The only example is early work on *Drosophila*, which mostly focused on late stages of PGS patterning and not on their early formation (29), but showed clearly that the interactions among the segment-polarity genes in the anterior segments are different from those in other segments. Nonetheless, the non-canonical expression of segment-polarity genes in the anteriormost segments has been shown in many arthropod species, although it has not always been pointed out explicitly. Expression of *en* in the PGS appears later than expected based on their position in myriapods (37-39), in spiders (40) and in crustaceans (41, 42). Stripe splitting of the early *hh* stripe in the PGS has been shown in spiders (25), in myriapods (23, 26) and in insects (27, 28). Expression of *wg* in early patterning of the anterior segments has, been studied in a few insect species, and has been shown to have an early “head blob” expression in the ocular segment, similar to what we have shown (31). Despite the patchy nature of the data, the phylogenetic spread of the evidence suggests very strongly that there are numerous differences in determination and patterning between the PGS and segments posterior to them in all arthropods and these differences are ancient and conserved.

### The PGS are not serially homologous to the trunk segments

Arthropod segments are said to be serially homologous structures. The exact definition of serial homology has been debated since the early days of comparative morphology (34, 43). While the standard definition of homologous structures is that they are descendent from the same structure in a common ancestor (6), structures within the same organism obviously do not have an ancestor. Thus, we must turn to other definitions of homology. The emerging paradigm within evolutionary developmental biology sees homologous structures as being patterned by conserved gene regulatory networks (44). This definition works for serial homology as well, since several structures in a single organism can be patterned by the same GRN. Wagner (34) defines serial homology as: “two body parts of the same organisms are serially homologous if they result from the repeated activation of the same character identity network” (p. 418). Similarly, Tomoyasu et al. (45) identified serially homologous structures as being orchestrated by the same developmental system. The conserved GRN (Wagner’s character identity network) underlying serially homologous segments is the segment-polarity gene network. This is the most conserved aspect of the segmentation cascade and is repeatedly activated in the formation of all segments in all arthropods studied to date (2, 9). We have shown that the three anterior segments of the arthropod head, collectively known as the pre-gnathal segments, do not share this conserved GRN. The various components of the segment-polarity network are expressed at different relative times and in different relative positions, and have different functional interactions. We therefore assert that under the definition of serial homology given above, the PGS are not serially homologous to the other segments in the arthropod body. Since many of the unusual aspects of the PGS are shared among different arthropod clades, we believe they are homologous within arthropods, an idea supported by neuroanatomy, by the expression of Hox genes and by the fossil record (19).

### Implication for the evolution of the arthropod head

The three anterior segments seem to be a distinct set of segments in terms of both evolutionary history and development. A segmented body with an anterior head tagma predates the common ancestor of arthropods (1, 46-48). The earliest branching stem group arthropods probably had a head made up of a single segment. Fossils with preserved nervous tissue of the stem arthropods *Kerygmachela* (49) and *Lyrarapax* (50) indicate that these animals had a single brain ganglion, corresponding to procephalon, as in the brain of tardigrades and thus likely primitive for Panarthropoda as a whole (51, 52). The upper-stem arthropod *Fuxianhuia* already has a tri-partite brain, as evident from exceptionally preserved fossils of this species (53). Most lower-stem arthropods had a single raptorial appendage pair, located on the protocerebral segment. Upper-stem arthropods (which unite with crown group arthropods in a grouping collectively known as Deuteropoda (54)) have their raptorial appendages on the second or deutocerebral segment (Fig. 6).

**Figure 6.**
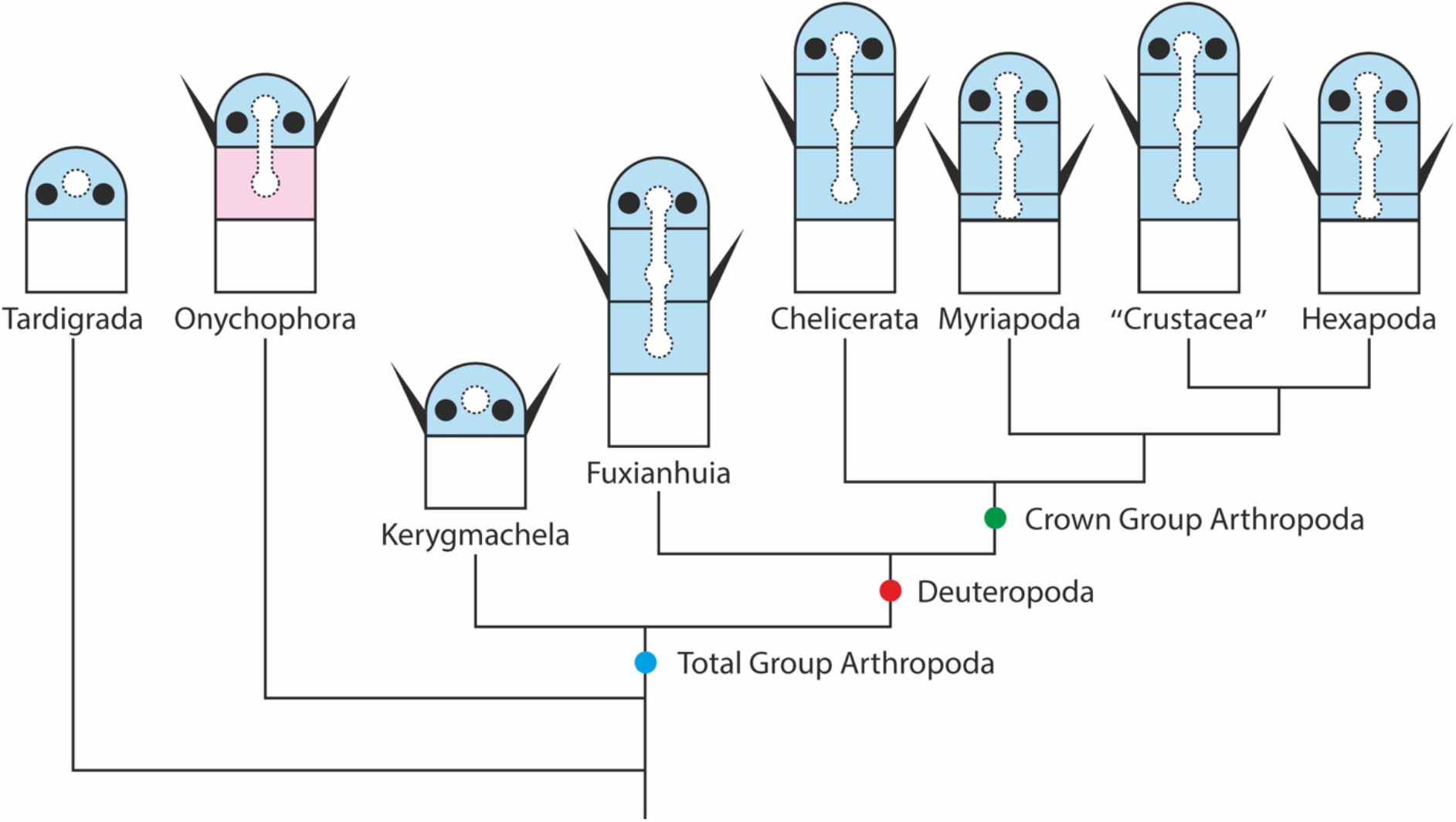
The change in head structure mapped on a phylogenetic tree of Panarthropoda. Each clade is represented by a scheme of the procephalon, marked in blue, and the segment posterior to it, marked in white. Dotted shape – the dorsal ganglion/ganglia. Black circles – eyes. Lateral black spikes – the anterior-most paired appendages. The third (tritocerebral) segments is shown to be reduced in Myriapoda and Hexapoda, where it becomes the appendageless intercalary segment. The Onychophoran second segment is shown in pink, to indicate that it is probably not homologous to the deutocerebral segment in Deuteropoda. The tree and the head structures are based on (19)

The transition between a single segmented head with a protocerebral appendage to a three-segmented head with a deutocerebral appendage is one of the most dramatic changes in early arthropod history and the most difficult to reconcile with developmental biology (55). It has been suggested that this transition occurred by the recruitment of two additional trunk segments to the head (19). If this scenario were true, we would expect at least the two “new” head segments to be serially homologous to the trunk segments, which is evidently not the case. We suggest a novel scenario for the evolutionary transition between a single-segmented head in lower stem arthropods and a three-segmented head in Deuteropoda, involving the splitting of an ancient head unit into three.

We suggest that in the early history of arthropods the body was made up of a series of homonomous trunk segments, with a single unit making up the head and a single neuropil functioning as a brain. This head was patterned and differentiated via a separate developmental pathway than the trunk segments, using some of the same genes that were used in the trunk segments, but with different interactions. As the brain expanded, it split into three parts, concomitantly splitting the surrounding morphological structures into three distinct units. The developmental mechanism through which this was achieved was a three-fold spatial repetition of the process generating the head, so that it generated three segmental units.

When a gene undergoes duplication, often each new copy takes on part of the roles originally carried out by the parent gene, a phenomenon known as sub-functionalization. By analogy, we suggest that when the ancestral head split to become a three-segmented head, each of the new segments took up some of the structures and functions of the ancestral head. This suggestion provides a possible solution to the debate regarding the homology of the deutocerebral raptorial appendage of Deuteropoda and the frontal raptorial appendage of lower stem arthropods such as *Kerygmachela*. When the ancestral single head segment split, the second of the resulting segments inherited the raptorial appendage of the original segment. The two appendages can thus be seen to be homologous, despite their different segmental position.

Indeed, we can say the three PGS are not serial homologs of each other, but rather – adopting once again the terminology from gene evolution – paralogs of each other. After splitting, they continued to evolve independently, free from the constraints of a shared gene regulatory network. This is consistent with the differences in the specifics of gene expression among the different segments, as we have shown here. The degradation of the tritocerebral segment to a rudimentary intercalary segment led to reduced and late expression of several of the segment-polarity genes in the insect intercalary segment. This process occurred convergently in myriapods, but with different molecular consequences.

## Conclusions

We have shown that the gene regulatory network patterning the pre-gnathal segments in the insect *Oncopeltus* is fundamentally different from that patterning all other segments. Patchy, but phylogenetically broad data from other arthropods indicates that this is a general phenomenon. We conclude that the three PGS have an evolutionary history that is independent from trunk segments and suggest they evolved through triplication of an ancestral single-segment head. With this new insight, it should be possible to reinterpret the change in the morphology of the head throughout arthropod evolution, as represented in the fossil record. This insight also opens the door for more detailed analyses of the development of the head in extant arthropods with the aim of reconstructing the precise changes in developmental regulation that lead to the evolution of the complex head we see today.

## Materials and Methods

### Animal culture and egg collection

*O. fasciatus* cultures are kept in a temperature-controlled room at 25°C in plexiglass boxes. each box contains several dozens of insects. To collect eggs, which are available year-round, cotton balls are placed in a box with sexually mature bugs, until a female lays a clutch on them or placed there for a pre-determined window of time. Eggs are kept in a 25°C incubator until they reach the desired stage of development.

### Antisense Digoxigenin-labeled probes and dsRNA preparations

The primers for the antisense Digoxigenin (DIG) labeled RNA probes and for the dsRNA for parental RNA interference were designed from the published *O. fasciatus* genome (NCBI accession number: PRJNA229125) for all relevant genes using the bioinformatics software Geneious (56).

Both probes and dsRNA were made using Sigma-Aldrich T7 RNA polymerase and buffer. Probes were made with Sigma-Aldrich Digoxigenin-labeled ribonucleotides (RNA). dsRNA was made using ribonucleotides from Lucigen.

### RNA interference using dsRNA parental injections

Virgin females are sedated with CO2 gas and injected with about 3-5 μL of dsRNA at an average concentration of 2 (+/- 0.5) mg/mL of respective mRNA in the ventral abdomen. The injected females recover overnight. The following day, each female is introduced to 1-2 adult males and placed in a box with cotton balls. Egg collecting starts from the third day of egg-laying.

### eggs fixation and peeling

*O. fasciatus* eggs are submerged in tap water and boiled for three minutes and then immediately placed on ice for at least 5 minutes. The water is then replaced with a 50%/50% mixture of 12% paraformaldehyde in PBS-T (phosphate buffered saline + tween) / heptane and put in a shaking wheel for primary fixation. All liquids are then removed, followed by several washes with 100% methanol. The embryos can be kept at −20°C. To prepare for in situ hybridizations, *O. fasciatus* eggs are peeled and post-fixed in 4% paraformaldehyde in PBS-T for 90-120 minutes in a rotating plate.

### In situ hybridization staining

Fixed embryos are stepwise transferred to PBS-T solution and placed in a hybridization buffer for at least 1 hour in 60°C. Antisense probes (Digoxigenin labeled RNA) are added to the buffer for a final concentration of 1mg/1mL probe/hybridization solution, and incubated at 60°C overnight. The embryos are washed several times with hybridization buffer to remove excess probes, and then PBS-T before transferred to 10% normal goat serum/PBS-T (blocking) solution for epitope blocking. AP-conjugated antibodies against Digoxigenin are added to a final concentration of 1:2500 antibodies/blocking solution, and the embryos are placed at 4°C overnight.

The embryos are then washed with PBS-T and transferred to a staining solution. We stain the embryos until we observe noticeable staining or until the embryo gains a purple tint, suggesting non-specific staining. We transfer the embryos stepwise to 70% glycerol/PBS-T and keep them at 4°C.

### Microscopy

*O. fasciatus* pictures were taken using an AZ100 stereoscope with Nikon Digital Sight DS-Fi1 camera.

## Acknowledgements

This work was funded by a grant from the Israel Science Foundation # 120/16. We thank Greg Edgecombe for discussions and for critical comments on an earlier version of this manuscript.

## Supplementary figures

**Supplementary Figure 1.**
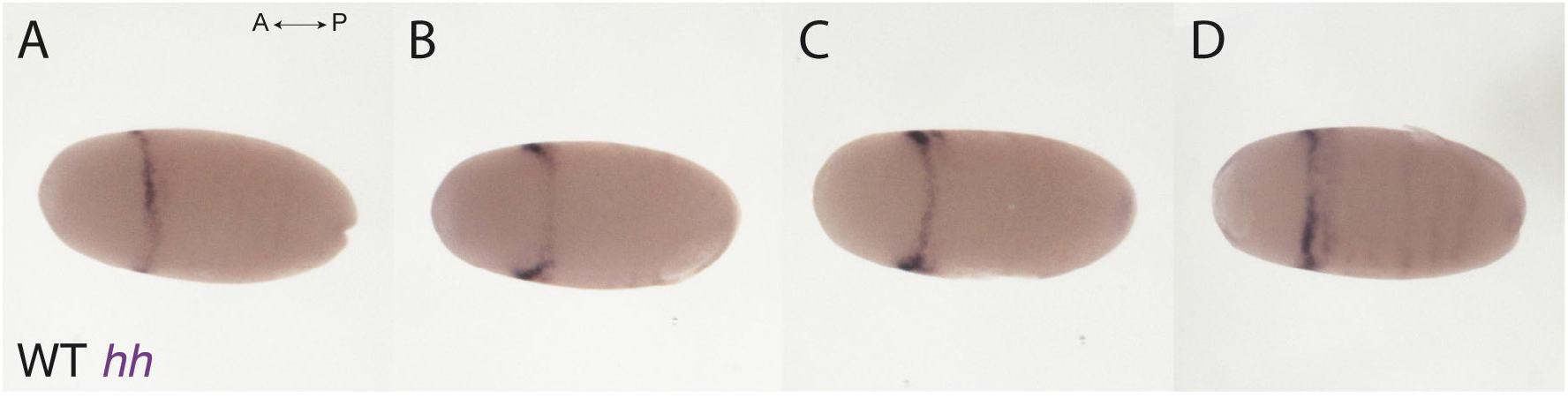
The development of wildtype expression of *hedgehog* in blastoderm embryos. A-D) from left to right: embryos from ~30 hours after egg laying (hAEL) to ~36 hAEL. an anterior ring of *hh* expression appears ~30 hAEL, which splits gradually, resulting in two stripes correlating to the ocular and antennal segments. the splitting continues during invagination, which starts at ~36 hAEL.

**Supplementary Figure 2.**
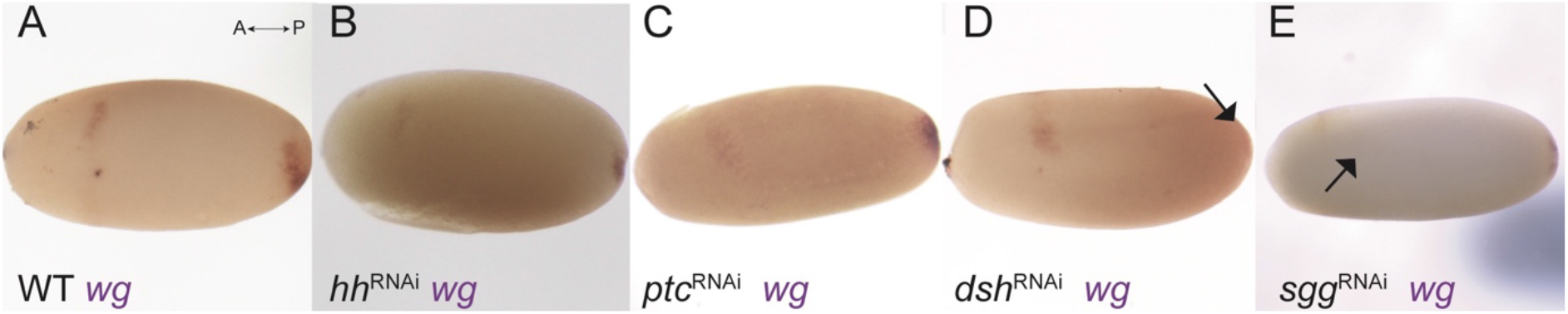
Expression of *wingless* in embryos ~32 hours after egg laying (hAEL) following pRNAi against different genes. (A) wildtype expression of *wg* is in an anterior patch which will be the ocular staining (“head blob”), and a posterior patch marking the future site of invagination. (B) Expression of *wg* in *hh*-RNAi embryos shows no significant difference from wildtype. (C) Expression of *wg* in *ptc*-RNAi embryos shows no significant difference from wildtype. (D) Expression of *wg* in *dsh*-RNAi embryos is missing in the posterior future site of invagination (black arrow), the anterior patch is the same as in wildtype embryos. (E) Expression of *wingless* in *sgg*-RNAi embryos is missing the anterior ocular staining (black arrow), the posterior patch is the same as in wildtype embryos.

